# Microtubules tune mechanosensitive cell responses

**DOI:** 10.1101/2020.07.22.205203

**Authors:** Shailaja Seetharaman, Benoit Vianay, Vanessa Roca, Chiara De Pascalis, Batiste Boëda, Florent Dingli, Damarys Loew, Stéphane Vassilopoulos, Manuel Théry, Sandrine Etienne-Manneville

## Abstract

Mechanotransduction is a process by which cells sense the mechanical properties of their surrounding environment and adapt accordingly to perform cellular functions such as adhesion, migration and differentiation. Integrin-mediated focal adhesions are major sites of mechanotransduction and their connection with the actomyosin network is crucial for mechanosensing as well as the generation and transmission of forces onto the substrate. Despite having emerged as major regulators of cell adhesion and migration, the contribution of microtubules to mechanotransduction still remains elusive. Here, we show that actomyosin-dependent mechanosensing of substrate rigidity controls microtubule acetylation, a tubulin post-translational modification, by promoting the recruitment of the alpha-tubulin acetyl transferase (αTAT1) to focal adhesions. Microtubule acetylation, in turn, promotes GEF-H1 mediated RhoA activation, actomyosin contractility and traction forces. Our results reveal a fundamental crosstalk between microtubules and actin in mechanotransduction, which contributes to mechanosensitive cell adhesion and migration.

## Main

Cells sense the physical properties of their environment, translate them into biochemical signals and adapt their behaviour accordingly. This process known as mechanotransduction is crucial during development as well as in the adult during physiological and pathological conditions such as cell migration, wound healing and cancer^1,2^. Integrin-mediated focal adhesions (FAs) sense the matrix rigidity, control the generation of actomyosin-dependent forces and the transmission of these traction forces onto the substrate, as well as contribute to mechanosensitive cell responses such as migration^3,4^. In addition to the actin cytoskeleton, microtubules are also key regulators of 2D and 3D cell migration^5-8^. Several studies have demonstrated the role of the actomyosin cytoskeleton and FAs in mechanotransduction, however, very little is known about microtubules in this context. In this study, we used astrocytes, whose migration is highly dependent on efficient microtubule dynamics, to address the role of microtubules in rigidity sensing and mechanosensitive migration^9-11^.

One of the crucial factors affecting the functions of the microtubule network is post-translational modifications (PTMs) of tubulin such as acetylation, which occurs at the K40 residue of α-tubulin. The enzyme responsible for microtubule acetylation, αTAT1 (α-tubulin acetyltransferase 1, also called as MEC-17), is present in the lumen of microtubules ^12^ and is highly specific to α-tubulin K40 (Fig. S1A). On the other hand, the enzymes involved in deacetylation at K40 are histone deacetylase family member 6 (HDAC6) and sirtuin type 2 (Sirt2) (Fig. supplementary S1A), both of which target other substrates as welln ^13^. We have previously shown that microtubule acetylation promotes FA turnover and cell migration ^11^. Thus, we investigated whether the extracellular matrix rigidity affects microtubule acetylation. Astrocytes were plated sparsely on polyacrylamide hydrogels of different rigidities: 1.26 kPa, 2 kPa, 9 kPa and 48 kPa (Fig. 1A and supplementary S1A). Astrocytes on soft substrates exhibited lower levels of acetylated tubulin than cells on stiff substrates as evidenced by a lower ratio of acetylated tubulin to the total tubulin in case of the former (Fig. 1A and supplementary S1A). Subsequently, to determine whether microtubule acetylation may be triggered by a mechanism involving adhesion and spreading, we plated cells on adhesive micropatterns (area 2500 μm^2^) printed on 2 kPa and 40 kPa hydrogels. Similar to stiff substrates, cells on soft substrates adopted a crossbow shape and identical spread area, and yet, microtubule acetylation was increased on stiff substrates as compared to softer substrates (Fig. supplementary S1B), In contrast, tubulin detyrosination, another PTM, was not affected by increased substrate rigidity (Fig. supplementary S1C), and had no effect on astrocyte adhesion and migration 11, suggesting that rigidity sensing specifically affects microtubule acetylation. Thus, we focused on understanding the role of microtubule acetylation in response to matrix rigidity.

**Figure 1:**
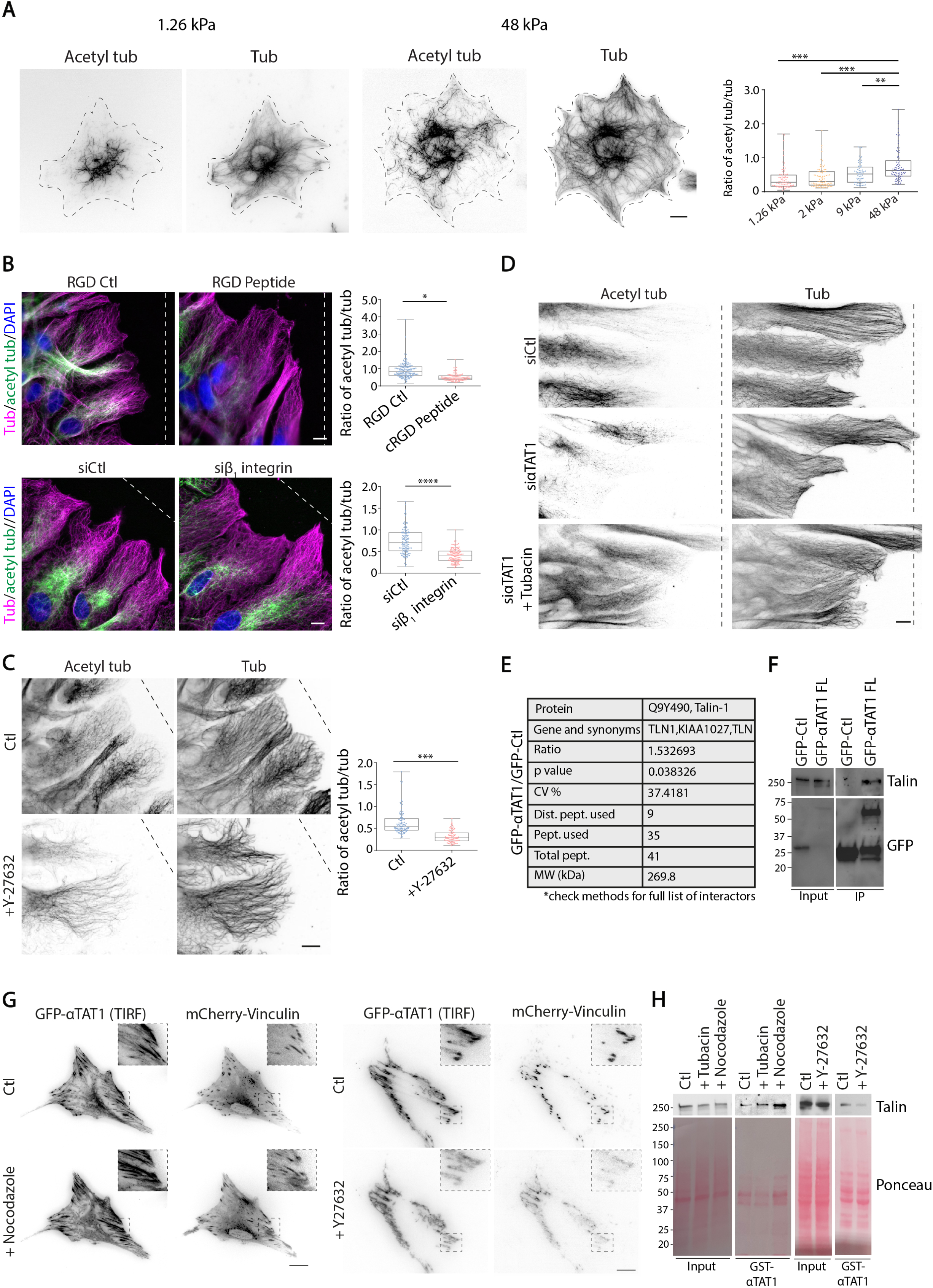
Integrin-mediated signalling and substrate rigidity regulate microtubule acetylation. **A**. Inverted-contrast epifluorescence images of astrocytes plated on polyacrylamide gels of different rigidities (1.26 kPa, 48 kPa), stained with acetylated tubulin and α-tubulin. Graph shows the ratio of the intensities of acetylated tubulin over total tubulin intensity of each cell; n > 67 cells, N = 4 independent experiments; one-way ANOVA followed by Tukey’s multiple comparison’s test. **B**. Astrocytes were treated with RGD control or RGD peptides (upper panels) prior to wounding or transfected with siCtl or siβ_1_ integrin (lower panels), allowed to migrate for 8 h. Images show migrating astrocytes stained with acetylated tubulin, α-tubulin and DAPI. Graph shows the ratio of the intensities of acetylated tubulin over total tubulin of each cell; n ≥ 100 cells, N = 3 independent experiments; one-way ANOVA followed by Tukey’s multiple comparison’s test. **C**. Inverted-contrast epifluorescence images of astrocytes, treated with DMSO (Ctl) or ROCK inhibitor Y-27632 for 2 h, and stained with acetylated tubulin and α-tubulin. Graph shows the ratio of the intensities of acetylated tubulin over total tubulin intensity of each cell; n ≥ 80 cells, N = 3 independent experiments; Student’s t-test. **D**. Inverted-contrast epifluorescence images of migrating astrocytes transfected with siCtl or siαTAT1 and treated with or without Tubacin prior to wounding. Representative images from N > 3 independent experiments are shown. **E**. Table shows the mass spectrometry data obtained on Talin-1 following the analysis of αTAT1 interactors. **F**. Immunoprecipitations using anti-GFP nanobodies were performed with lysates from astrocytes transfected with GFP-Ctl or GFP-αTAT1. Samples were analysed by immunoblotting using talin and GFP antibodies. Representative blot from N = 3 independent experiments is shown. **G**. TIRF images of GFP-αTAT1 and epifluorescence images of mCherry-vinculin expressing astrocytes before and after nocodazole or Y27 treatment. Insets represent regions of the cell where αTAT1 and vinculin colocalise. Representative images from N = 3 independent experiments. **H**. Pulldowns using GST-αTAT1 resin were performed with lysates from astrocytes treated with tubacin, nocodazole or Y27632, and analysed by red Ponceau staining and immunoblotting using talin. Representative blots from N = 3 independent experiments are shown. Scale bar (A-D): 10 μm, (G): 20 μm; *p<0.05, **p<0.01, ***p<0.001, ****p<0.0001.

Cells sense the matrix rigidity and trigger a cascade of signalling pathways downstream of integrins. To determine the role of integrin signalling in controlling microtubule acetylation, we used a scratch induced migration assay to trigger integrin activation at the wound edge^9^. Addition of cyclic RGD (cRGD) peptide, to prevent the binding of integrins to the RGD motif of extracellular matrix proteins, reduced tubulin acetylation (Fig. 1B). Furthermore, depletion of β_1_ integrin using a siRNA (Fig. supplementary S1D) also resulted in a significant decrease in tubulin acetylation as compared to control cells (Fig. 1B). The enzymes responsible for microtubule acetylation and deacetylation are αTAT1 and HDAC6 respectively. Thus, we used previously characterized siRNAs targeting αTAT1 (Fig. supplementary S1A) to decrease acetylation and Tubacin^11,14^, a drug which increases microtubule acetylation by inhibiting HDAC6 (Fig. supplementary S1A), without modifying the acetylation of other HDAC6 substrates such as histones^14,15^. On depleting αTAT1, we observed that αTAT1 was responsible for integrin-mediated microtubule acetylation (Fig. 1C). These results suggest that β_1_ integrin signalling promotes microtubule acetylation on rigid substrates.

Mechanosensing requires actomyosin contractility^16,17^. We treated cells plated on glass coverslips with the ROCK inhibitor, Y-27632, which strongly reduces actomyosin contractility^18^. Under this condition, tubulin acetylation is highly reduced (Fig. 1D and supplementary S1E), strongly suggesting that mechanosensing at FAs controls microtubule acetylation.

The recruitment and/or activation of αTAT1 remains unknown^19^. Therefore, we carried out a quantitative mass spectrometry screen to identify interacting partners of αTAT1 using HEK cells. The mass spectrometry data (data available via ProteomeXchange with identifier PXD015871; check methods for details; Fig. supplementary S1F) revealed interesting potential interactors, amongst which the proteins enriched in the gene ontology for focal adhesions are depicted in red (Fig. supplementary S1F). One of the significant interactors on the mass spectrometry screen was talin (Fig. 1E), a mechanosensitive partner of integrins. The interaction of αTAT1 with talin was further confirmed by co-immunoprecipitation in astrocytes (Fig. 1F). In addition, using TIRF microscopy, we observed that GFP-αTAT1 strongly localised at mCherry-vinculin-positive FAs (Fig. 1G)^11^. Since αTAT1 is also present within microtubules (Fig. supplementary S1G)^11,12^, we tested whether the recruitment of αTAT1 to FAs was dependent on microtubules. To address this, GFP-αTAT1 and mCherry-vinculin expressing astrocytes were treated with nocodazole after acquiring a short movie of GFP-αTAT1 localisation at FAs. Nocodazole-treated cells displayed larger adhesions as observed in other cell types (Fig. 1G, movie 1)^20,21^ and higher levels of αTAT1 at FAs (Fig. 1G, movie 1). This was also verified using GST-αTAT1 pulldowns, wherein, the interaction of αTAT1 with talin increased in nocodazole or tubacin-treated astrocytes compared to control (Fig. 1H). Similarly, by TIRF microscopy as well as GST-αTAT1 pulldowns, we observe that with the loss of FAs upon Y-27632 treatment, αTAT1 interaction with talin is diminished in agreement with decreased tubulin acetylation, suggesting a tension-dependent recruitment of αTAT1 at focal adhesions (Fig. 1G and 1H, movie 2). Altogether, these results show that mechanosensing at focal adhesions triggers the recruitment of αTAT1 to promote microtubule acetylation. It also suggests that the recruitment of αTAT1 may be a consequence of mechanosensitive changes in talin conformation^22^.

We then tested whether microtubule acetylation may be involved in mechanosensitive cell functions. First, we analysed the impact of substrate rigidity on the distribution of FAs by immunostaining of the FA-associated protein, paxillin^4,23^. Astrocytes were sparsely plated on polyacrylamide hydrogels of different rigidities: 1.26 kPa, 2 kPa, 9 kPa and 48 kPa (Fig. 2A). Quantification of the density of FAs (Fig. supplementary S2A) indicates that cells on soft substrates display FAs throughout the cell surface, whereas, FAs were predominantly seen at the periphery in cells adhering to stiffer substrates (Fig. 2A). Thus, changes in substrate rigidity alter FA distribution in astrocytes, allowing us to assess the impact of microtubule acetylation in this phenomenon. Following αTAT1 depletion (Fig. supplementary S1A), FAs were distributed throughout the cell surface independently of the rigidity of the substrate, i.e., αTAT1-depleted cells on 48 kPa had a similar distribution of FAs as control/siαTAT1 cells on 1.26 kPa substrates (Fig. 2B, 2D and supplementary S2B). To confirm that the changes in FA distribution act through tubulin acetylation, we treated cells plated on different rigidities with tubacin. Tubacin did not have an effect on the FA localisation in cells plated on 48 kPa substrates (Fig. 2C, 2D and supplementary S2B). In contrast to αTAT1 depletion, tubacin treatment in cells plated on 1.26 kPa substrates mimicked the phenotype (FAs at the cell periphery) observed in control/tubacin-treated cells plated on stiff matrices (Fig. 2C, 2D and supplementary S2B). Since cell spreading on different substrate rigidities can have an effect on the distribution of FAs, we therefore plated αTAT1-depleted cells on micropatterned hydrogels to observe FAs in cells of similar spread area. In line with our prior results, cells on soft substrates or αTAT1-depleted cells on stiffer substrates displayed FAs throughout the cell surface (Fig. supplementary S2C). Together with previous findings that microtubule acetylation controls membrane vesicle delivery at FAs and FA dynamics^11,24^, these results show that the rigidity-sensing dependent microtubule acetylation controls the mechanosensitive distribution of FAs (Fig. 2E).

**Figure 2:**
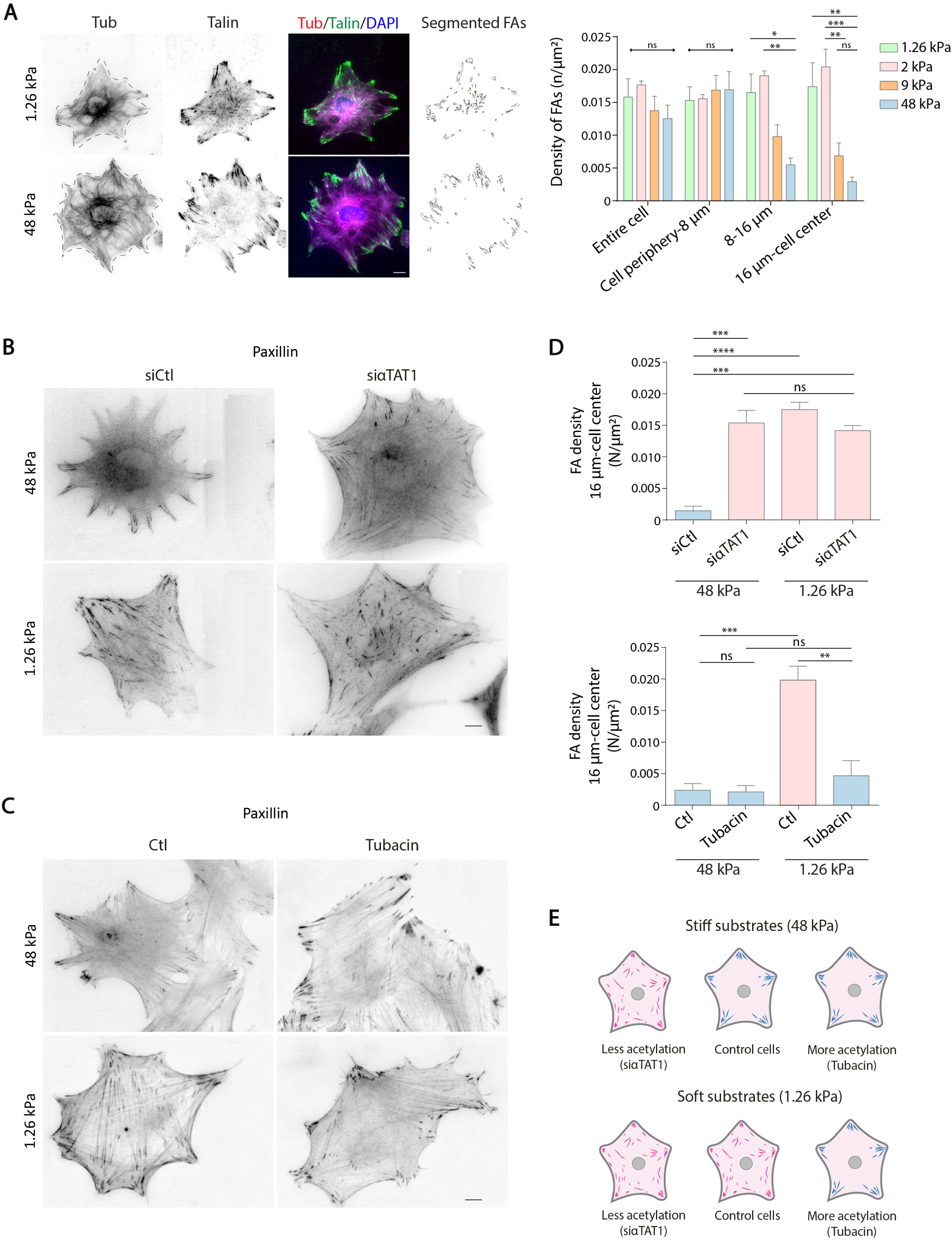
Microtubule acetylation tunes mechanosensitivity of focal adhesions (FAs). **A**. Inverted epifluorescence images of astrocytes plated on polyacrylamide gels of different rigidities (1.26 kPa and 48 kPa), stained with α-tubulin and talin. Images shown correspond to the same cells depicted in Fig. 1A. Talin images were segmented by adjusting the threshold to detect FAs. Histogram shows the mean ± SEM of FA density (number of FAs/μm^2^) in different regions of the cells plated on substrate of indicated rigidities; n = 40 for 1.26 kPa, 51 for 2 kPa, 94 for 9 kPa, 77 for 48 kPa, N = 3 independent experiments; two-way ANOVA followed by Tukey’s multiple comparison’s test. **B**. Inverted epifluorescence images of astrocytes transfected with siCtl and siαTAT1, plated on different substrate rigidities and stained with paxillin. **C**. Inverted epifluorescence images of astrocytes treated with with Niltubacin and tubacin, plated on different substrate rigidities and stained with paxillin. **D**. Histogram shows the mean ± SEM of FA density (number of FAs/μm^2^) in the central region (16μm-cell center) of cells depicted in B and C; n = 60 for 1.26 kPa siCtl, 36 for 1.26 kPa siαTAT1, 54 for 48 kPa siLuc and 47 for 48 kPa siαTAT1; n = 59 for 1.26 kPa Niltubacin, 55 for 1.26 kPa tubacin, 46 for 48 kPa Niltubacin and 37 for 48 kPa tubacin; N = 3 independent experiments; one-way ANOVA followed by Tukey’s multiple comparison’s test. **E**. Schematic showing the effects of microtubule acetylation on the distribution of FAs in cells plated on polyacrylamide gels of different rigidities. Scale bar (A-C): 10 μm; *p<0.05, **p<0.01, ***p<0.001, ****p<0.0001.

Next, we investigated the impact of αTAT1 depletion on the FA-associated cytoskeleton. In migrating astrocytes, the actin network comprises of longitudinal stress fibres connected to FAs at the front of the leader cells as well as interjunctional transverse arcs connecting neighbouring cells at adherens junctions (Fig. 3A)^18,25^. In αTAT1-depleted cells, the transverse arcs of actin were dramatically reduced and the longitudinal fibres did not extend to the front of migrating cells (Fig. 3A and supplementary S2D). Associated with these longitudinal fibres, FAs were located further back in the protrusion rather than at the front of leader cells (Fig. 3A)^11^. Myosin light chain phosphorylation (pMLC) in control cells is predominantly seen at the leading edge of migrating cells, however, in αTAT1-depleted cells, pMLC was barely visible at the leading edge of cells and was only associated with the remaining actin fibres at the cell center, similar to myosin IIa distribution (Fig. 3A and 3B). Moreover, intermediate filaments (visualised using vimentin), which play a major role in regulating FAs and collective migration of astrocytes ^25^ and normally extend from the perinuclear region to the cell periphery close to FAs^25,26^, were noticeably absent from the front of αTAT1-depleted cells and frequently appeared fragmented (Fig. 3C). We then looked closely at the effect of αTAT1 on the cytoskeletal organisation at FAs by using platinum-replica transmission electron microscopy (EM) on unroofed migrating astrocytes located at the wound egde. As by light microscopy, EM images showed that FAs connected to actin bundles were distributed further within the protrusion in case of siαTAT1 cells as compared to a highly organised and parallel set of FAs at the leading edge of control cells (Fig. 3D-1, 3D-2 and 3D-3). From the high magnification views of FAs in the control cells, microtubules were often seen along actin cables reaching FAs (Fig. 3D-1i, marked with white arrows). Intermediate filaments were also clearly visible, intertwined with the actin filaments at FAs (Fig. 3D-1ii and iii, marked with yellow arrows). In siαTAT1 cells, the actin bundles near FAs were strikingly thinner than those in controls (Fig. 3D-1ii, 3D-2iv and Fig. 3D-3v). We consistently observed a lack of microtubules and intermediate filaments associated with FAs in αTAT1-depleted cells (Fig. 3D-2iv and 3D-3v). All these results strongly support a role for αTAT1 in the cytoskeletal organisation at FAs.

**Figure 3:**
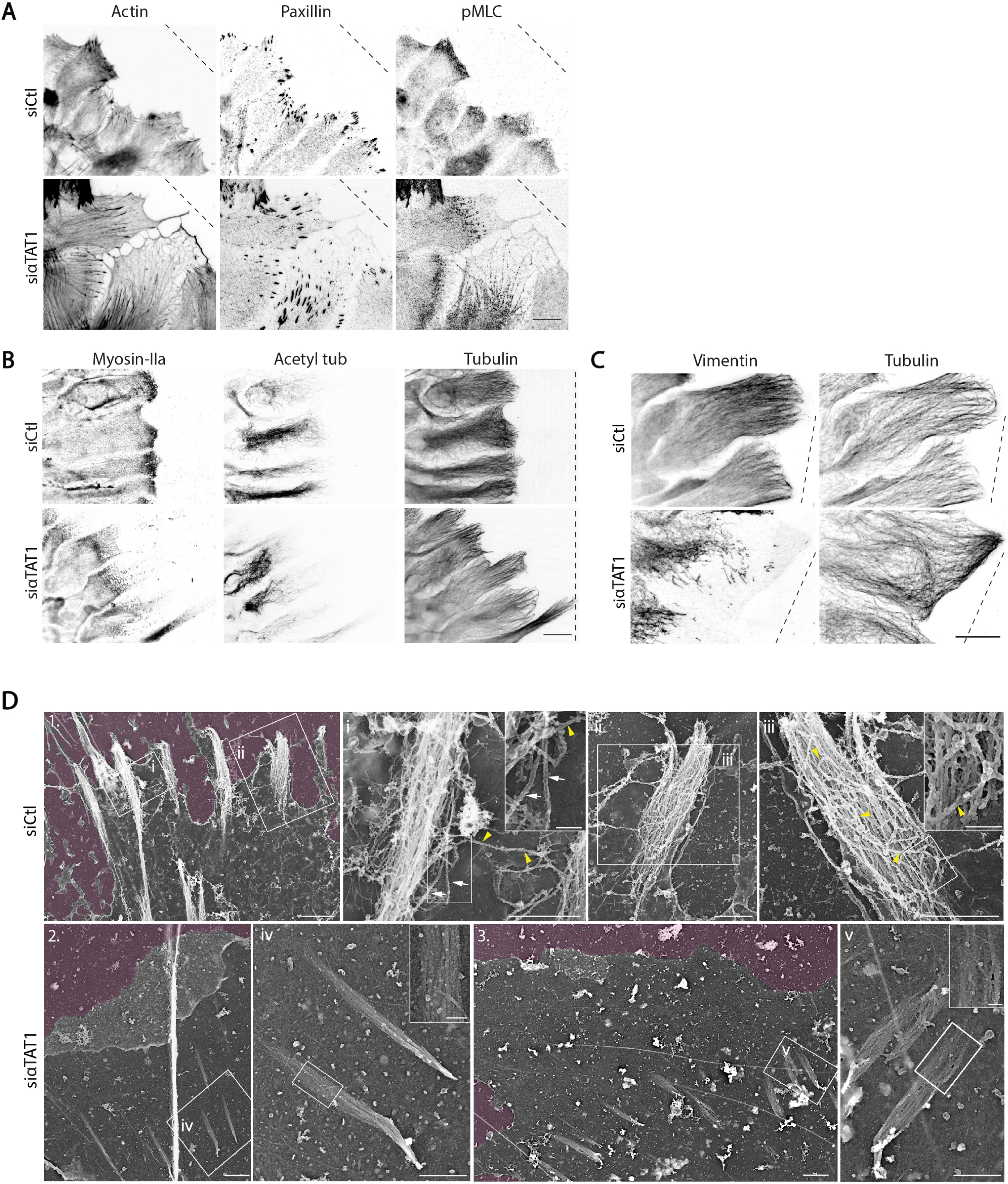
Microtubule acetylation reorganises the actomyosin and intermediate filament networks. **A-C** Inverted epifluorescence images of astrocytes transfected with siCtl or siαTAT1 and stained with (**A**) phalloidin, pMLC and paxillin; (**B**) myosin IIa, acetylated tubulin and α-tubulin; and (**C**) vimentin and α-tubulin. Images are representative of N = 3 independent experiments. Scale bar (A-C): 20 μm. **D**. Ultrastructural organization of focal adhesion-associated cytoskeleton in siCtl or siαTAT1 depleted cells. (1) Platinum replica electron microscopy (PREM) survey view of the cytoplasmic surface of the leading edge in siCtl unroofed cells. Boxed regions correspond to focal adhesions. Extracellular space is pseudo-coloured in purple. (i, ii) High magnification views corresponding to the boxed regions in panel 1. White arrows indicate microtubules and yellow arrowheads denote intermediate filaments. (iii) Zoom-in region corresponding to the boxed region in marked in region ii. Scale bar: 1 µm. White arrows indicate microtubules and yellow arrowheads denote intermediate filaments. Scale bar: 2 µm and 1 µm. (2, 3) PREM survey view of the cytoplasmic surface of the leading edge in αTAT1-depleted cells. Extracellular space is pseudo-coloured in purple. Scale bars: 10 µm, 1 µm (inset). (iv, v) High magnification views corresponding to the boxed regions in panel 2 and 3 respectively. Scale bar: 1 µm.

One major impact of mechanosensing is the adaptation of traction forces to the rigidity of the substrate^27^. We thus asked whether microtubule acetylation might affect traction forces by plating control or αTAT1-depleted astrocytes on crossbow-shaped micropatterned hydrogels. By traction force microscopy, we observed that αTAT1 depletion resulted in lower traction force production on 40 kPa substrates (Fig. 4A, 4B and supplementary S3A). In contrast, overexpression of GFP-αTAT1 increased traction energies and forces and also rescued the effect observed on αTAT1 knockdown (siαTAT1 + GFP-αTAT1; Fig. 4A, 4B, supplementary S3B and S3D) when cells were plated on 40 kPa. In addition, tubacin-treated cells on softer 2 kPa hydrogels showed increased traction energies and forces, comparable to control cells on 40 kPa (Fig. 4A, 4C and supplementary S3C). Thus, the level of microtubule acetylation dictates traction forces exerted on the substrate through FAs. This further illustrates the essential role of the mechanosensitive regulation of microtubules in force transmission.

**Figure 4:**
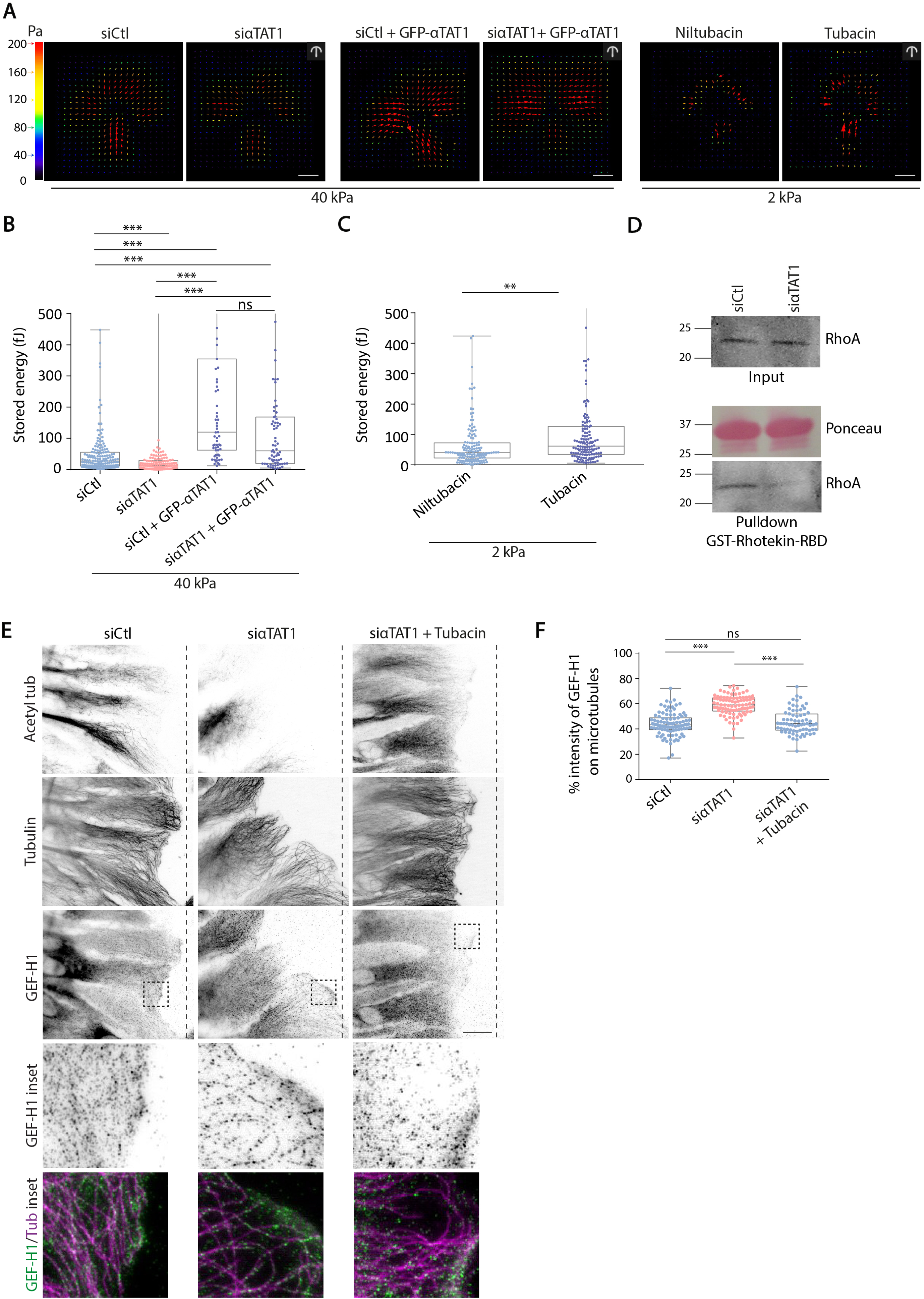
Microtubule acetylation promotes traction force generation and alters GEF-H1 localisation. **A**. Stress-field maps of astrocytes transfected with siCtl, siαTAT1, siαTAT1 + GFP-αTAT1, treated with Niltubacin or Tubacin, and plated on crossbow-shaped micropatterned polyacrylamide gels of 40 kPa or 2 kPa. **B, C**. Corresponding stored energies (in Joules, J) of cells in each of the above mentioned conditions. Values represent mean ± SEM stored energies of cells within the range of 0 to 5 × 10^−13^ J; n ≥ 142 cells for siCtl and siαTAT1, n ≥ 71 cells for siCtl + GFP-αTAT1 and siαTAT1 + GFP-αTAT1, n ≥ 137 cells for Niltubacin and tubacin, N = 3 independent experiments; One-way ANOVA followed by Tukey’s multiple comparison’s test (graph B) or Student’s t-test (graph C). **D**. GST-Rhotekin pulldowns were performed using siCtl or siαTAT1-transfected astrocytes. Red Ponceau and anti-RhoA western blot analysis of lysates. Representative blot from N = 3 independent experiments is shown. **E**. Inverted epifluorescence images of migrating astrocytes transfected with siCtl, siαTAT1 and siαTAT1 treated with tubacin, stained with acetylated tubulin, α-tubulin, GEF-H1. Graph represents mean ± SEM of the percentage of GEF-H1 colocalised with microtubules; n ≥ 73 cells; N = 3 independent experiments; One-way ANOVA followed by Tukey’s multiple comparison’s test or Student’s t-test. Scale bar (A): 10 μm, (E) 20 μm; **p<0.01,b ***p<0.001, ns – not significant.

The crucial role of microtubule acetylation in the controlling cytoskeletal organisation and traction forces led us to further investigate the molecular mechanisms involved in this process. We focused on RhoA, a small G protein of the Rho family, well-known for promoting stress fibre formation and actomyosin contractility, via its effector ROCK and MLC phosphorylation. Pulldown of GTP-bound active RhoA using GST-Rhotekin beads showed that αTAT1 depletion reduced RhoA activity (Fig. 4D), suggesting that microtubule acetylation may promote actomyosin contractility by activating RhoA. RhoA activation is mediated by Guanine nucleotide exchange factors (GEFs)^28^. Amongst these GEFs, GEF-H1 (also known as ARHGEF2) is a microtubule-bound RhoGEF which, when released from microtubules triggers the Rho-ROCK signalling cascade and cell contractility^29,30^. Previously, substrate stiffness was suggested to correlate with GEF-H1 activity and actomyosin contractility^31^. In addition, GEF-H1 was recently shown to be controlled by the interaction of microtubules with integrin-mediated adhesions^32^, leading us to investigate whether integrin-mediated microtubule acetylation could affect the association of GEF-H1 with microtubules. In control astrocytes plated on rigid glass coverslips, GEF-H1 localised partially (approximately 42%) on microtubules but also as dots in the cytosol, which are known to depict the active/released GEF-H1 (Fig. 4E and 4F)^33^. In contrast, in αTAT1-depleted cells, GEF-H1 was found predominantly localised on microtubules (approximately 60%; Fig. 4E and 4F). Tubacin treatment of αTAT1-depleted cells led to GEF-H1 release into the cytosol rescuing the effect of αTAT1 depletion and confirming the role of microtubule acetylation in the GEF-H1 localisation (Fig. 4E and 4F). Altogether, these results show that microtubule acetylation promotes the release of GEF-H1 from microtubules into the cytoplasm, and strongly suggest that rigidity-dependent microtubule acetylation contributes to mechanotransduction by enabling RhoA activation and actomyosin contractility.

The involvement of microtubule acetylation in mechanotransduction and in the mechanosensitive regulation of FAs and acto-myosin contractility led us to investigate its influence on cell migration, which has often been described as a mechanosensitive cellular response^2,34^. To this end, we developed a collective migration assay on hydrogels, where microdropping a small amount of a chemical (sodium hydroxide) induced a circular wound in the cell monolayer (Fig. 5A, movie 3). Wild-type astrocytes migrated significantly slower on 2 kPa gels than on 48 kPa gels (Fig. 5B, movie 4), implying that astrocyte migration speed is affected by substrate rigidity. Most importantly, αTAT1 depletion abolished the increase of cell speed observed on stiff 48 kPa gels (Fig. 5C, movie 5), where the cell migration speed of αTAT1-depleted cells plated on 48 kPa was similar to that of control/αTAT1-depleted cells on 2 kPa substrates. Thus, we demonstrate that microtubule acetylation is required for mechanosensitive regulation of astrocyte collective migration.

**Figure 5:**
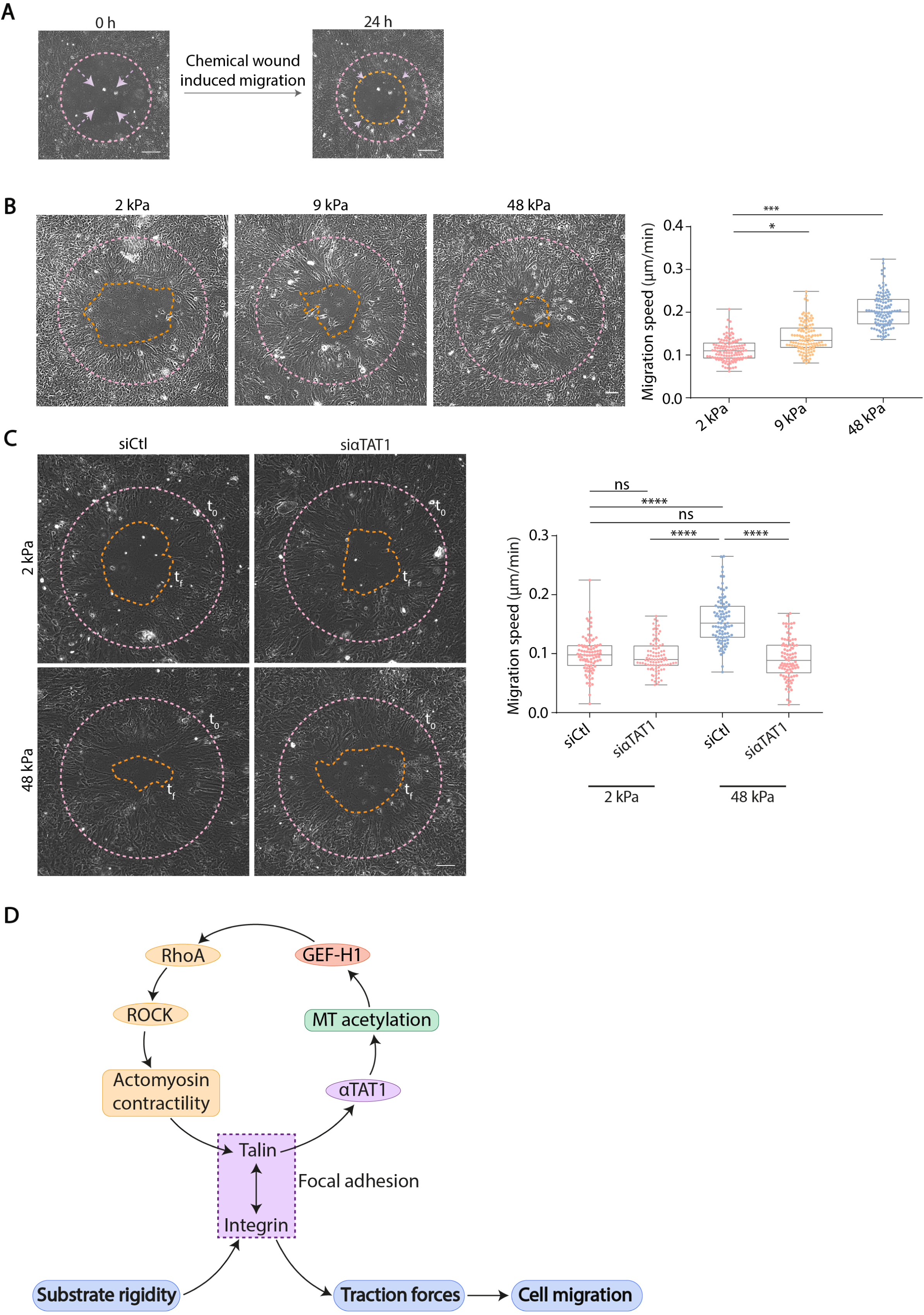
Microtubule acetylation is required for mechanosensitive migration. **A**. Phase contrast images showing a chemical wound migration assay. **B**. Phase contrast images showing astrocyte migration 24 h after wounding in chemical wound assays performed on 2 kPa, 9 kPa and 48 kPa substrates. Pink dotted lines represent initial wound edge and orange dotted lines represent the final wound edge. Graph represents the migration speed (in μm/min) of cells at the wound edge; n ≥ 115 cells, N = 3 independent experiments; one-way ANOVA followed by Tukey’s multiple comparison’s test. **C**. Phase contrast images showing the migration siCtl or siαTAT1-transfected astrocytes 8h after wounding in chemical wound assays performed on 2 kPa and 48 kPa substrates. Pink dotted lines represents initial wound edge and orange dotted lines represent final wound edge. Graph represents the migration speed (in μm/min) of cells at the wound edge; n ≥ 92 cells, N = 3 independent experiments; one-way ANOVA followed by Tukey’s multiple comparison’s test. Scale bar (B, C): 100 μm; ****p<0.0001, ns – not significant, *p<0.05, ***p<0.001. **C**. Proposed working model. Actomyosin-dependent matrix rigidity sensing through integrin and talin promotes αTAT1 interaction with talin, and microtubule (MT) acetylation. MT acetylation facilitates the release of GEF-H1 from MTs, which activates RhoA, triggers the Rho/ROCK signalling and increases actomyosin contractility, traction forces at focal adhesions and collective cell migration.

Our results show that microtubules are regulated in response to substrate rigidity sensing and in turn, play a key role in mechanotransduction by participating in mechanosensitive cellular responses (Fig. 5D). Downstream of integrin-mediated signalling, mechanosensing controls the recruitment of αTAT1 to FAs and induces microtubule acetylation. Microtubule acetylation tunes mechanosensitive distribution of FAs and force generation, thereby, contributing to cellular responses to substrate rigidity. We hypothesize that the actomyosin-sensitive association of αTAT1 with talin might be crucial in transmitting signals to the cytoskeleton, leading to microtubule acetylation upon integrin activation (or wounding). In agreement with our findings, formins, which control actin dynamics, have also been shown to facilitate microtubule acetylation^35,36^. How αTAT1 enters the lumen of microtubules still remains unclear although one can speculate that αTAT1 accesses the lumen through microtubule lattice defects or through the open ends^19^. Growing microtubule ends are often seen in close proximity to FAs at the leading edge of migrating cells^8^. It is highly plausible that part of the pool of αTAT1, at FAs, enters the lumen through these microtubule open ends in the vicinity of adhesions.

We show that microtubule acetylation reorganises the actomyosin network and promotes traction forces. Therefore, we propose a feedback mechanism involving a crosstalk between microtubules and actin wherein, actomyosin-dependent mechanosensing promotes microtubule acetylation which, in turn, facilitates the release of GEF-H1 from microtubules into the cytosol to increase RhoA activity, cell contractility and traction forces. It was recently shown that uncoupling microtubules from FAs results in a similar release of microtubule-bound GEF-H1 into the cytosol^32^, which then triggers myosin IIA assembly and increased cell contractility through RhoA. Suppression of RhoA activity in the absence of αTAT1 might be due to the sequestering of GEF-H1 by non-acetylated microtubules. Whether microtubule acetylation directly or indirectly induces the release of GEF-H1 remains unclear. One can speculate that changes in the conformation of the microtubule lattice due to intraluminal acetylation of tubulin may facilitate the release of GEF-H1 from microtubules. Alternatively, increased acetylation which makes microtubules more resilient^37,38^ may promote softening, bending or curving of microtubules, thereby, facilitates the release of GEF-H1 from microtubules. Interestingly, microtubule acetylation also affects the association of intermediate filaments with actin bundles at FAs. Since intermediate filaments have also been involved in the control of FA dynamics, actomyosin contractility as well as GEF-H1 activity^25,33^, the microtubule-actin interplay described here may also involve intermediate filaments, whose role in mechanotransduction is still elusive.

In response to substrate rigidity sensing, cells perform essential functions such as migration^39^. In the absence of αTAT1, cells plated on stiff substrates produce less traction forces and in turn, migrate slower. This can be considered counter-intuitive for single cell migration where traction forces appear to correlate with migration speed, however, collective cell migration relies on the transmission of forces between the leaders and followers. Cell-cell junctions not only transmit forces between cells but help maintain the integrity of the monolayer^18,40^, which improves collective and directed cell migration. Alteration of microtubule acetylation did not induce any detachment of leader cells from followers and nor were there any effects in directionality or persistence during migration of αTAT1-depleted cells^11^. Therefore, we propose a model in which, on stiff substrates, increased microtubule acetylation would trigger higher traction forces in leader cells, which would transmit these pulling forces to followers and increase collective migration speed.

In conclusion, our results depict a crosstalk between the actin and microtubule cytoskeletal networks (Fig. 5D), whereby microtubule acetylation, downstream of rigidity-dependent integrin-mediated signalling, alters actomyosin contractility as well as focal adhesion distribution and dynamics to promote mechanosensitive migration of astrocytes, thus closing a crucial feedback loop governing mechanotransduction at FAs (Fig. 5D).

## Methods

### Cell culture

Primary astrocytes were obtained from E17 rat embryos^10^. Use of these animals is in compliance with ethical regulations and has been approved from the Prefecture de Police and Direction départementale des services vétérinaires de Paris. Astrocytes were grown in 1g/L glucose DMEM supplemented with 10% FBS (Invitrogen, Carlsbad, CA), 1% penicillin-streptomycin (Gibco) and 1% Amphotericin B (Gibco) at 5% CO_2_ and 37°C.

### Cell nucleofection

Astrocytes were transfected with Lonza glial transfection solution and electroporated with a Nucleofector machine (Lonza). Cells were then plated on appropriate supports previously coated with poly-L-Ornithine (Sigma). Experiments are carried out 3 or 4 days post-transfection and comparable protein silencing was observed. siRNAs were used at 1 nmol and DNA was used at 5 µg. siRNA sequences used were: Luciferase (control) UAAGGCUAUGAAGAGAUAC; αTAT1 rat (siαTAT1-1): 5’-ACCGACACGUUAUUUAUGU-3’ and 5’-UUCGAAACCUGCAGGAACG-3’; αTAT1 rat (siαTAT1-2): 5’-UAAUGGAUGUACUCAUUCA-3’ and 5’-UCAUGACUAUUGUAGAUGA-3’; β_1_ integrin (AUUGCCAGAUGGAGUAACA). Constructs used were: GFP-αTAT1 and GST-αTAT1 (gift from Philippe Chavrier, Institut Curie and Guillaume Montagnac, Institut Gustave Roussy) and mCherry-vinculin. siαTAT1-1 and siαTAT1-2 are pools of two siRNAs each. For all experiments, siαTAT1-2 has been used due to better knockdown efficiency as seen in Fig. S1A and previously characterized in ^11^.

### Cell treatment

Cells were plated on appropriate supports and allowed to grow for 2-3 days after nucleofection. 2 mM Tubacin (Sigma) or 2 mM Niltubacin (negative control for Tubacin; Enzo Life Sciences) were added prior to wounding cells. Similarly, RGD peptide (Enzo Life Sciences) and RGE Control peptide (Enzo Life Sciences) were added prior to wounding. ROCK inhibitor, Y-27632, was added 6 h after wounding and 2 h before fixation. For TIRF experiments, 10 μM nocodazole or Y-27632 were added after 10 min of acquisition. For pull-downs, 10 μM nocodazole or Y-27632 were added 1 h before cell lysis.

### Preparation of homogenously coated polyacrylamide hydrogels

Protocol to prepare polyacrylamide hydrogels was adapted from ^34,41,42^. Coverslips were plasma cleaned for 3 min and silanised for 10 min using a solution of 1% (v/v) 3-(trimethoxysilyl)propyl methacrylate and 1% (v/v) acetic acid in ethanol. Coverslips were then washed twice with absolute ethanol and dried. A solution of polyacrylamide (proportions of acrylamide and bisacrylamide in the solution define the rigidity of the hydrogel) was prepared. 2.5 µl 10% APS and 0.25 µl TEMED was added and mixed well with the solution. A 50 µl drop of the solution was added over each coverslip (20×20 mm) and immediately, an 18×18 mm coverslip was placed gently over the solution. The solution was allowed to polymerise for 1 h at room temperature. HEPES was added over the coverslips to detach the top glass. The polymerised gel was then activated under UV for 5 min using Sulpho-SANPAH and washed with HEPES twice. The hydrogels were then coated with 100 µg/ml of rat tail collagen I overnight at 4°C. The excess collagen was washed once with PBS and approximately 5 × 10^4^ cells/ml were plated on hydrogels.

### Micropatterning

Coverslips were plasma-cleaned for 45 s and incubated with 0.1 mg/ml poly-l-lysine/polyethylene glycol (PLL-PEG) diluted in 10 mM HEPES for 30 min at room temperature. Excess of PLL-PEG was allowed to slide down by gravity, and the coverslips were dried and stored at 4°C overnight before printing. Micropatterns were printed on previously prepared PLL-PEG coverslips for 3 min with specifically designed chrome masks and coated with 50:50 v/v fibronectin/fibrinogen mixture (20 μg/ml each) diluted in fresh NaHCO_3_, pH 8.3, 100 mM, for 30 min at room temperature. Micropatterned coverslips were washed thrice in NaHCO_3_ and used immediately for transfer on hydrogels. Plated cells (approximately 6 × 10^4^ cells/ml) were allowed to adhere for 16 h before imaging/fixation. Crossbow patterns have an area of ∼2500 µm^2^.

### Traction force experiments on micropatterned substrates

After printing and coating micropatterned coverslips, gel mixtures were prepared as follows: 40.4 kPa – 100 µl 40% acrylamide, 120 µl 2% bisacrylamide, 280 µl water (final concentration of 8% acrylamide and 0.48% bisacrylamide in water); 2.61 kPa – 50 µl 40% acrylamide, 25 µl 2% bisacrylamide, 425 µl water (final concentration of 8% acrylamide and 0.048% bisacrylamide in water). 5 µl of fluorescent microbeads (FluoSpheres; Molecular Probes) were incubated with 0.1 mg/ml PLL-PEG on a rotator for 1 h at 4°C prior to mixing with the gel mixture. The PLL-PEG coated beads were washed and centrifuged thrice at 1000 rpm for 10 min with 10 mM HEPES. These beads were then mixed with 165 µl of the gel mixture. After adding 1 µl APS and 1 µl TEMED, the solution was added as a 25 µl drop on a silanised 20×20 mm coverslip. The protein-coated micropatterned coverslip was then gently placed on the gel mixture. The solution was allowed to polymerize for 25 min at room temperature. After detaching the micropatterned glass from the polymerized gel, 6 × 10^4^ cells/ml were plated on the gels and allowed to adhere and spread for 16 h before TFM experiments. Stacks of single micropatterned cells were acquired before and after trypsin treatment. Acquisitions were performed with a Nikon Eclipse Ti-E epifluorescence inverted microscope, with a 40× 1.49 NA air objective equipped with a pco.edge sCMOS camera and Metamorph software. Cells were maintained at 5% CO_2_ and 37°C in normal astrocyte medium during acquisition.

### Migration assays

For in vitro wound healing assays, cells were plated on appropriate supports (dishes, plates, coverslips or glass-bottom MatTek). Cells were allowed to grow to confluence and fresh medium was added the day prior to the experiment. The cell monolayer was scratched with a p200 pipette tip to induce migration and imaged immediately/8 h post wounding.

For migration on soft substrates, cells were plated on hydrogels of different rigidities in 6-well glass bottom plates (MatTek) and allowed to grow for 72 h after transfection before creating a chemical wound. 0.05 M NaOH was gently dropped on the cells using a microinjector. The dead cells and debris were washed off gently using PBS. Fresh medium was added to the cells. Cells were placed on the microscope for live imaging.

Acquisition was started 30 min after wounding. Movies were acquired with a Zeiss Axiovert 200M or a Nikon Eclipse Ti-E epifluorescence inverted microscope with cells maintained at 5% CO_2_ and 37°C. All images were acquired with dry objective 10X 0.45 NA and an EMCCD camera/pco.edge sCMOS camera and Metamorph software. Images were acquired every 15 min for 24 h. Nuclei of leader cells were manually tracked with Fiji software (Manual Tracking plugin).

### Immunostaining

Cells were fixed with cold methanol for 3 min at -20°C, or 4% warm PFA, or 4% PFA + 0.2% glutaraldehyde + 0.25% Triton X-100 for 10 min at 37°C. After fixation with PFA, glutaraldehyde and triton, free aldehyde groups were quenched with a solution of 1 mg/ml sodium borohydride (Sigma-Aldrich) freshly added to cytoskeletal buffer (10 mM MOPS, 150 mM NaCl, 5 mM EGTA, 5 mM MgCl_2_, 5 mM glucose, pH 6.1) for 10 min on ice. Cells were permeabilised for 5 min with 0.1% Triton in case of PFA fixation. Coverslips were blocked for 1 h with 5% BSA in PBS. The same solution was used for primary and secondary antibody incubation for 1 h. Nuclei were stained with DAPI and coverslips were mounted with Prolong Gold.

Antibody anti-acetylated tubulin (clone 6-11B-1, mouse monoclonal, Sigma Aldrich), anti-Poly-Glu tubulin (1:1000, AbC0101, ValBiotech), anti-α-tubulin (MCA77G, rat; Biorad), anti-paxillin (610051, mouse monoclonal, lot 5246880; BD; and ab32084, rabbit monoclonal, clone Y133 and lot GR215998-1; Abcam), anti-talin (T3287, mouse monoclonal, clone 8D4, lot 035M4805V; Sigma-Aldrich), anti-GEF-H1 (ab155785, Abcam), anti-Phalloidin (176759, lot GR278180-3; Abcam), anti-vimentin (V6630, mouse monoclonal lot 10M4831; Sigma-Aldrich), anti-pMLC (3675S, S19, mouse, Cell signalling), anti-myosin IIA (non-muscle, M8064, rabbit polyclonal, Sigma-Aldrich). Secondary antibodies were Alexa Fluor 488 donkey anti–rabbit (711-545-152), Rhodamine (TRITC) donkey anti–rabbit (711-025-152), Alexa Fluor 647 donkey anti–rabbit (711-695-152), Alexa Fluor 488 donkey anti–mouse (715-545-151), Rhodamine (TRITC) donkey anti– mouse (715-025-151), Alexa Fluor 647 donkey anti-rat (711-605-152), and Alexa Fluor 488 donkey anti– rat (712-545-153); all from Jackson ImmunoResearch. Epifluorescence images were acquired with a Leica DM6000 microscope equipped with 40X 1.25 NA or 63X 1.4 NA objectives and recorded on a CCD camera with a Leica software.

### TIRF microscopy

Acquisitions were performed with a Nikon Eclipse Ti-E epifluorescence inverted microscope, with a 60× 1.49 NA oil objective equipped with a pco.edge sCMOS camera with Metamorph software. Cells were maintained at 5% CO_2_ and 37°C in normal astrocyte medium. After acquiring a 15 min movie of GFP-αTAT1 and mCherry-vinculin, nocodazole or Y27 were added and images were acquired every 2 min for 1 h.

### Transmission electron microscopy

Adherent astrocytes plated on glass coverslips were scratched to induce migration. 4 h after scratch, cells were disrupted by sonication as described previously ^43^. Coverslips were unroofed by scanning the coverslip with rapid sonicator pulses in KHMgE buffer (70 mM KCl, 30 mM HEPES, 5 mM MgCl_2_, 3 mM EGTA, pH 7.2). Paraformaldehyde 2%/glutaraldehyde 2%-fixed cells were further sequentially treated with 0.5% OsO_4_, 1% tannic acid and 1% uranyl acetate prior to graded ethanol dehydration and Hexamethyldisilazane substitution (HMDS, Sigma-Aldrich). Dried samples were then rotary-shadowed with 2 nm of platinum and 5-8 nm of carbon using an ACE600 high vacuum metal coater (Leica Microsystems). Platinum replicas were floated off the glass by 5% hydrofluoric acid, washed several times by floatation on distilled water, and picked up on 200 mesh formvar/carbon-coated EM grids. The grids were mounted in a eucentric side-entry goniometer stage of a transmission electron microscope operated at 80 kV (Philips, model CM120) and images were recorded with a Morada digital camera (Olympus). Images were processed in Adobe Photoshop to adjust brightness and contrast and presented in inverted contrast.

### Image analysis

Normalised mean intensity levels of acetylated and detyrosinated tubulin for immunofluorescence images were calculated as follows:

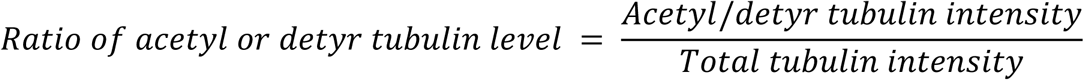

FA density was calculated as follows:

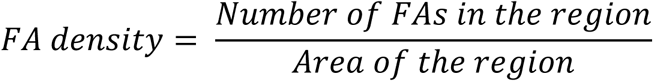

Different regions of cells (entire cell, cell periphery-8 μm, 8 μm-16 μm, 16 μm-cell center) were analysed as depicted in Fig. S2A.

For migration assays, nuclei of cells were manually tracked to determine the speed directionality and persistence of migration using the following formulae:

Mean velocity (*v*), persistance (*p*) and directionality (*d*) of cell migration are calculated as follows: for a given (*x, y*) coordinate of leading cell nucleus,

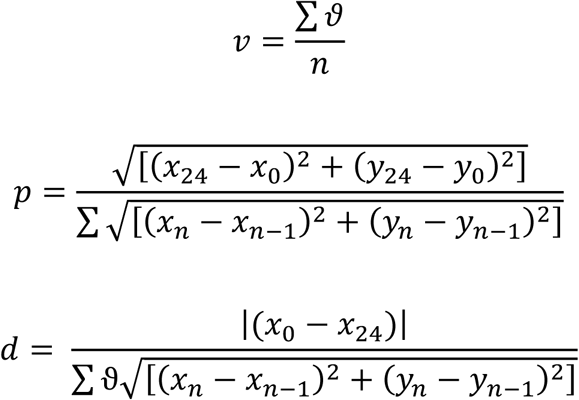

where *n* is the number of time points acquired and *ϑ* is cell velocity.

Analysis of TFM on micropatterns was performed with a custom-designed macro in Fiji based on work by ^44^. The topmost planes of beads before and after trypsinisation were selected and aligned using a normalized cross-correlation algorithm (Align Slices in Stack plugin). The displacement field was computed from bead movements using particle image velocimetry (PIV). Parameters for the PIV were three interrogation windows of 128, 64, and 32 pixels with a correlation of 0.60. Traction forces were calculated from the displacement field using Fourier transform traction cytometry and a Young’s modulus of 40 kPa or 2 kPa, a regularization factor of 10^−9^, and a Poisson ratio of 0.5.

For the localisation of GEF-H1 on microtubules, the percentage GEF-H1 on microtubules was calculated as follows:

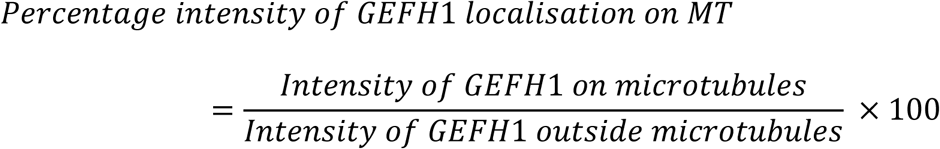

### Mass spectrometry

Proteins on beads were washed twice with 100 μL of 25 mM NH_4_HCO_3_ and we performed on-beads digestion with 0.2 μg of trypsine/LysC (Promega) for 1 h in 100 µL of 25 mM NH_4_HCO_3_. Sample were then loaded onto a homemade C18 StageTips for desalting. Peptides were eluted using 40/60 MeCN/H_2_O + 0.1% formic acid and vacuum concentrated to dryness. Online chromatography was performed with an RSLCnano system (Ultimate 3000, Thermo Scientific) coupled to an Orbitrap Fusion Tribrid mass spectrometer (Thermo Scientific). Peptides were trapped on a C18 column (75 μm inner diameter × 2 cm; nanoViper Acclaim PepMapTM 100, Thermo Scientific) with buffer A (2/98 MeCN/H_2_O in 0.1% formic acid) at a flow rate of 4.0 µL/min over 4 min. Separation was performed on a 50 cm x 75 μm C18 column (nanoViper Acclaim PepMapTM RSLC, 2 μm, 100Å, Thermo Scientific) regulated to a temperature of 55°C with a linear gradient of 5% to 25% buffer B (100% MeCN in 0.1% formic acid) at a flow rate of 300 nL/min over 100 min. Full-scan MS was acquired in the Orbitrap analyzer with a resolution set to 120,000 and ions from each full scan were HCD fragmented and analyzed in the linear ion trap.

For identification, the data was searched against the Homo sapiens (UP000005640) SwissProt database using Sequest^HF^ through proteome discoverer (version 2.2). Enzyme specificity was set to trypsin and a maximum of two missed cleavage site were allowed. Oxidized methionine, N-terminal acetylation, and carbamidomethyl cysteine were set as variable modifications. Maximum allowed mass deviation was set to 10 ppm for monoisotopic precursor ions and 0.6 Da for MS/MS peaks. The resulting files were further processed using myProMS ^45^ v3.6 (work in progress). FDR calculation used Percolator and was set to 1% at the peptide level for the whole study. The label free quantification was performed by peptide Extracted Ion Chromatograms (XICs) computed with MassChroQ version 2.2 ^46^. For protein quantification, XICs from proteotypic peptides shared between compared conditions (TopN matching) with no missed cleavages were used. Median and scale normalization was applied on the total signal to correct the XICs for each biological replicate. To estimate the significance of the change in protein abundance, a linear model (adjusted on peptides and biological replicates) was performed and p-values were adjusted with a Benjamini–Hochberg FDR procedure with a control threshold set to 0.05. Up-regulated proteins with at least 3 proteotypic peptides (fold change > 1.5 and p-value < 0.05) and the unique proteins identified only in the GFP-αTAT1 were used for gene ontology (GO) enrichment analysis by using GO::TermFinder tools (10.1093/bioinformatics/bth456) through myProMS.

The mass spectrometry proteomics data have been deposited to the ProteomeXchange Consortium via the PRIDE ^47,48^ partner repository with the dataset identifier PXD015871 (username; password: EC4DSdRf; project name: Potential interactors of alpha-tubulin acetyltransferase 1 (αTAT1/MEC17).

### Immunoprecipitations

Astrocytes were transiently transfected with pEGFP-c1 or GFP-αTAT1 using the calcium phosphate method. Cell lysates were prepared by scraping cells in lysis buffer 50 mM Tris pH 7.5, triton 20%, NP40 10%, 2 M NaCl with Complete protease inhibitor tablet (Roche, Indianapolis, IN) and centrifuged for 30 min at 13,000 rpm 4°C to pellet cell debris. Soluble detergent extracts were incubated with GST-GFP nanobody for 2 hr at 4°C prior to washing three times with wash buffer (50 mM Tris pH 7.5, 150 mM NaCl, 1 mM EDTA, 2.5 mM MgCl_2_). The resin was then mixed with Laemmli buffer and used for western blot and mass spectrometry analysis. HEK293 cells were used for mass spectrometry analysis in order to have enough amount of protein from cells expressing αTAT1.

### Rho activation assay

Rho activation assay was performed using a RhoA Pull-down Activation Assay Biochem Kit from Cytoskeleton Inc (BK036-S). In short, cells were lysed in ice-cold Cell Lysis Buffer plus 1x protease inhibitor cocktail. Cells were centrifuged at 10,000 g at 4°C for 10 min to pellet cell membranes and insoluble material. Part of the supernatant was stored as input for western blot. The remaining supernatant was divided equally into two parts (300-800 μg protein/tube). 1/15^th^ the volume of Loading Buffer was added to each tube (final conc. 15 mM). Then, 1/100^th^ the volume of GTPγS was added to one of the tubes and used as positive control (final conc. 0.2 mM). All tubes were incubated at room temperature for 15 min. The reaction was stopped by adding 1/10^th^ the volume of STOP Buffer to all tubes (final conc. 60 mM). Rhotekin-RBD (50 μg) beads were resuspended and added to the tubes. Tubes were rotated at 4°C for 1 h and centrifuged at 5000 g at 4°C for 3 min. Beads were washed with 500 μl each of Wash Buffer. 10-20 ul of Laemmli sample buffer was added to each tube and the bead samples were boiled for 2 min.

### Western blot

Cells lysates were obtained with Laemmli buffer composed of 60 mM Tris-HCl pH6.8, 10% glycerol, 2% SDS and 50 mM DTT with the addition of anti-protease (cOmplete cocktail, Roche 11 873 588 001). Samples were boiled 5 min at 95°C before loading on polyacrylamide gels. Transfer occurred at 100V for 1 h on nitrocellulose membranes. Membranes were blotted with TBST (0.2% Tween) and 5% milk and incubated 1 h with the primary antibody and 1 h with HRP-conjugated secondary antibody. Bands were revealed with ECL chemoluminescent substrate (Biorad).

Primary antibodies used: Antibody anti-acetylated tubulin (clone 6-11B-1, Sigma), α-tubulin (Biorad MCA77G), anti-β_1_ integrin (ab52971, Abcam), anti-GAPDH (MAB374, lot 2689153 Millipore), anti-talin (T3286 clone 8D4, lot 035M4805V, Sigma), anti-vinculin (lot 036M4797V, V9131, Sigma). Secondary HRP antibodies were all purchased from Jackson ImmunoResearch.

## Author contributions

S.S. designed, performed experiments, analysed results and wrote the paper; B.V. assisted in the setup of micropattern printing, TFM experiments and analysis, helped with data interpretation; V.R. performed immunoprecipitation and pull down experiments; C. D-P helped with experimental techniques and discussions; B. B optimised the GFP-nanobody conditions and IPs used for mass spectrometry sample preparation; F. D. carried out the MS experimental work; D. L. supervised MS and data analysis; S.V. performed EM experiments; M.T. provided ideas, helped with data interpretation and discussions; S. E-M. supervised the project, interpreted the results, and wrote the paper.

## Competing interests

Authors declare no competing interests.

## Supplementary figure legends

**Figure S1:**
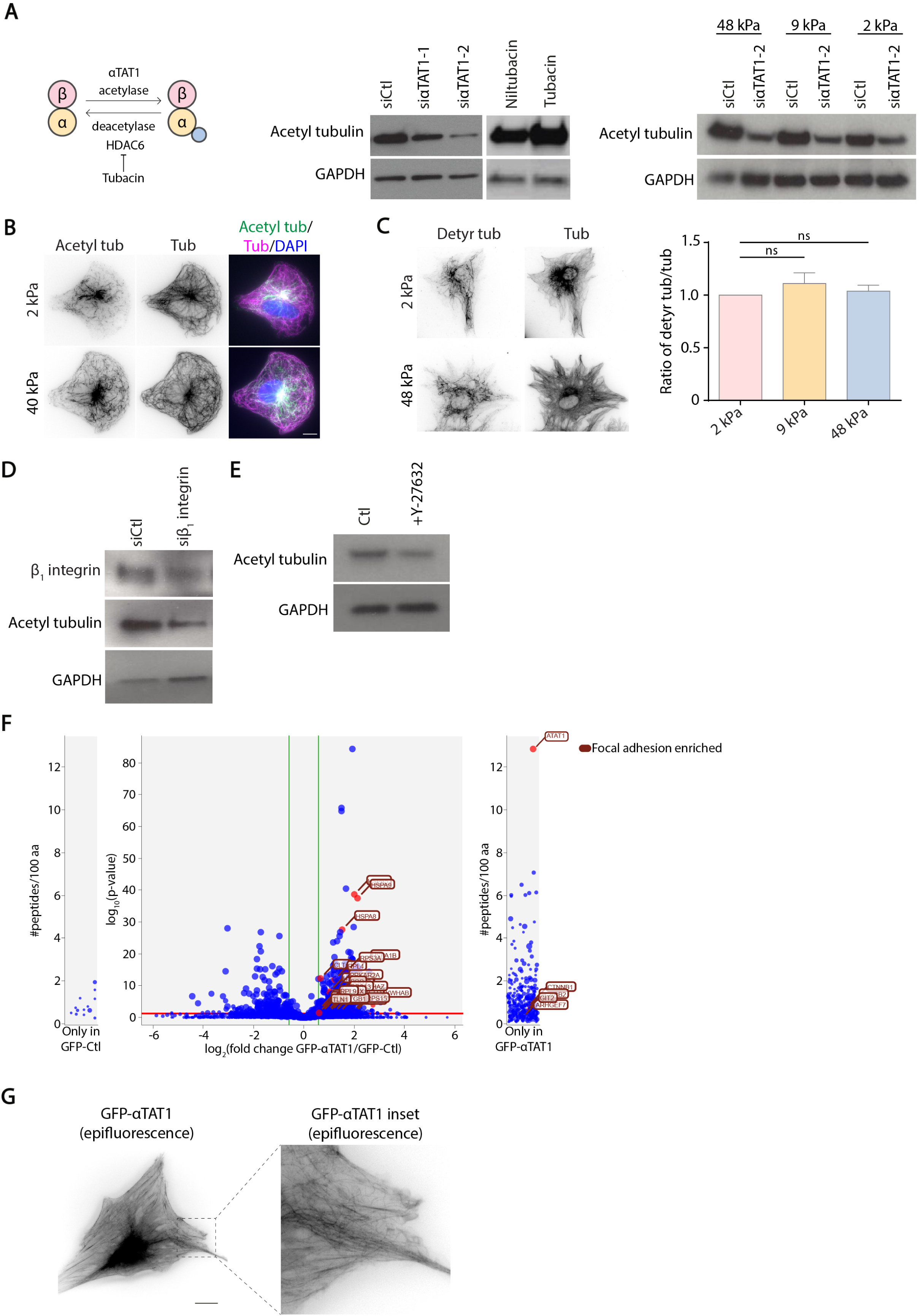
**A**. Schematic and western blots showing the tools used to manipulate the levels of microtubule acetylation in this study. 2 distinct sets of siRNAs targeting αTAT1 (siαTAT1-1 and siαTAT1-2) reduce microtubule acetylation and Tubacin, an inhibitor of HDAC6 (deacetylase), increases microtubule acetylation in astrocytes plates on hydrogels or glass. Representative blots from N = 3 independent experiments are shown. In the main figures, only results obtained with siαTAT1-2, which induces the strongest decrease in acetyl tubulin, are shown. Although less pronounced, siαTAT1-1 produced similar effects as siαTAT1-2, in experiments corresponding to Fig. 1D, 2B, 3A, 3B and 4E. **B**. Epifluorescence images of astrocytes transfected with indicated siRNAs and plated on crossbow-shaped micropatterned hydrogels of different rigidities, stained for acetylated tubulin and α-tubulin. **C**. Epifluorescence images showing detyrosinated tubulin and α-tubulin staining of astrocytes sparsely plated on hydrogels of different rigidities. Graph shows the intensity ratio of detyrosinated tubulin over total tubulin in each condition; N = 2 independent experiments; ns – not significant. **D**. Western blot showing the levels of β_1_ integrin, acetylated tubulin and GAPDH in astrocytes transfected with control siCtl or siβ_1_ integrin. **E**. Western blot showing the levels of acetylated tubulin and GAPDH in DMSO Ctl or Y-27632 treated (2 h) astrocytes, 6h after wound healing of the monolayer. **F**. Volcano plot analysis identifying interactors of αTAT1 in HEK cells. Binding partners were obtained by using quantitative label-free mass spectrometry analysis performed from four replicates. Shown are the fold changes (GFP-αTAT1/GFP-Ctl) of the quantified proteins with thresholds of 3 or more peptides, minimum absolute fold change of 1.5 (green lines) and maximum adjusted p-value of 0.05 (red line). Enriched protein interactors related to GO:0005925 focal adhesion (Ratio = 1.98 and p = 7.89 × 10^−5^) are shown (red). External plots show proteins with peptides identified only in one sample type (left in GFP-Ctl and right in GFP-αTAT1). The ratio of talin is indicated in the table. **G**. Epifluorescence images showing GFP-αTAT1 localisation on microtubules in astrocytes (as previously described in ^11^). Scale bar (B, C, G): 10 μm.

**Figure S2:**
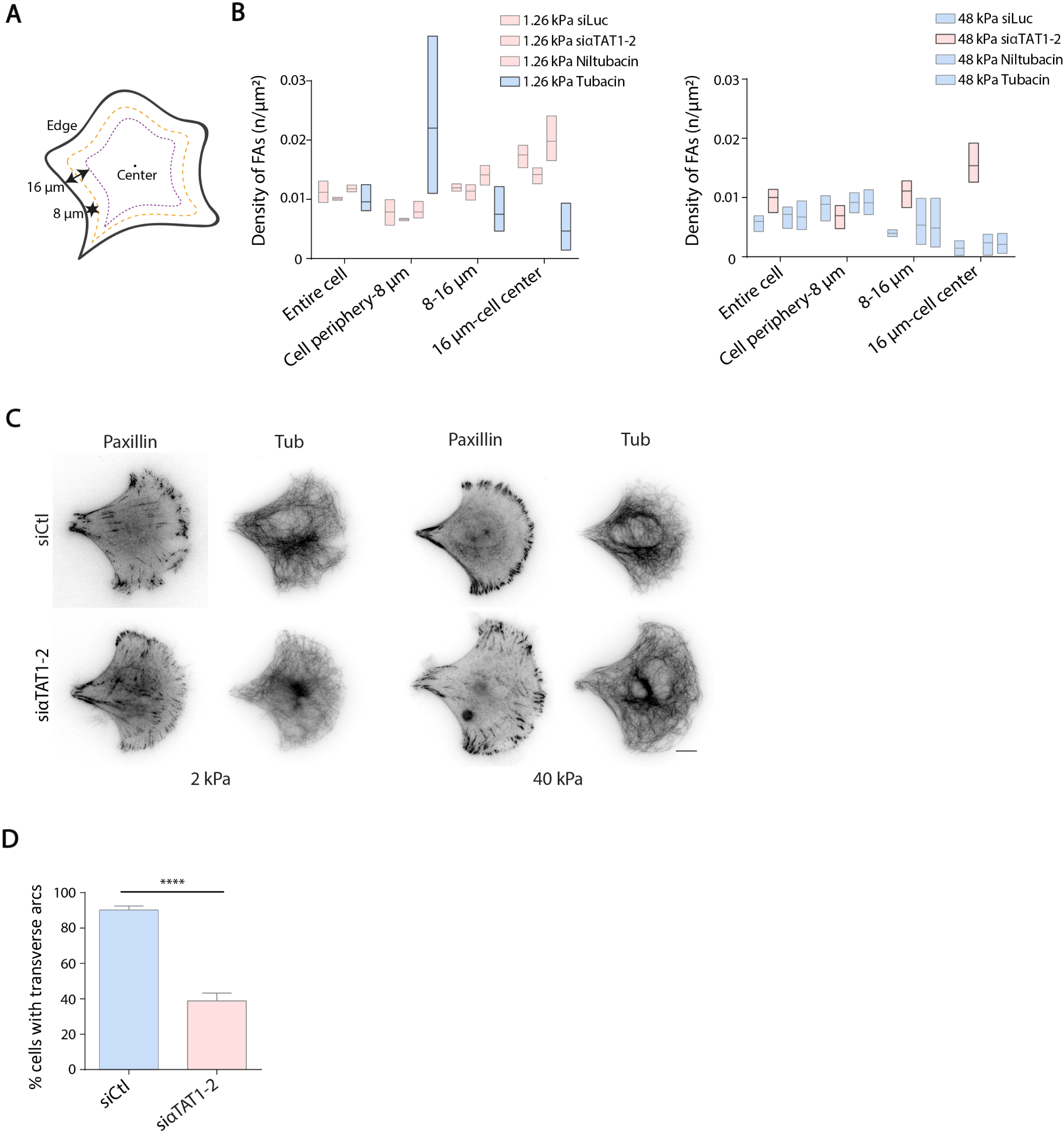
**A**. Schematic representation of the different cell regions used to quantify FA density (Fig. 2). **B**. Histograms show mean ± SEM of FA density (number of FAs/μm^2^) in different regions of astrocytes transfected with siCtl or siαTAT1 or treated with Niltubacin or Tubacin, and plated on 1.26kPa or 48 kPa substrates; N = 3 independent experiments. **C**. Inverted epifluorescence images of siCtl or siαTAT1-transfected astrocytes plated on crossbow-shaped micropatterned polyacrylamide gels of 40 kPa or 2 kPa, stained with paxillin and tubulin. Representative images from N = 3 independent experiments are shown. Scale bar: 10 μm **D**. Graph shows the percentage of siCtl or siαTAT1-transfected astrocytes with transverse interjunctional actin arcs; n ≥ 159 cells, N = 3 independent experiments; Student’s t-test; ****p<0.0001.

**Figure S3:**
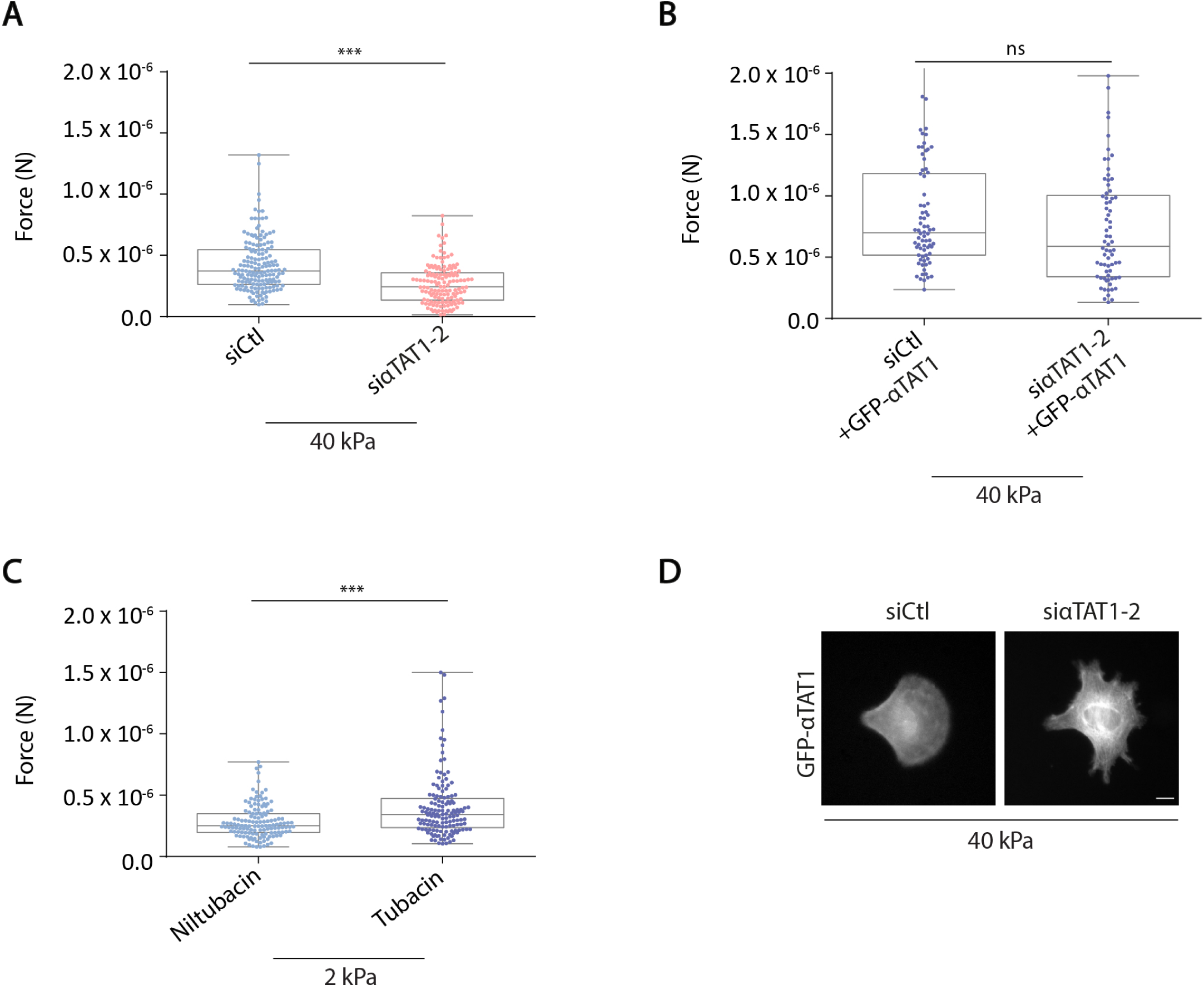
**A, B, C**. Traction forces in siCtl, siαTAT1, siCtl + GFP-αTAT1 or siαTAT1 + GFP-αTAT1 transfected cells, and Niltubacin or Tubacin treated cells are shown. Graphs represent forces (in N); n ≥ 142 cells for siCtl and siαTAT1, n ≥ 71 cells for siCtl + GFP-αTAT1 and siαTAT1 + GFP-αTAT1, n ≥ 137 cells for Niltubacin and tubacin, N = 3 independent experiments; Student’s t-test; ***p<0.001, ns – not significant. **D**. GFP-αTAT1 expression in cells transfected with siCtl + GFP-αTAT1 or siαTAT1 + GFP-αTAT1 is shown. Scale bar: 10 μm.

***Movie 1: GFP-αTAT1 localisation at FAs upon nocodazole treatment***. Cells were transfected with GFP-αTAT1 and mCherry-vinculin. First, images were acquired every 2 min for 15 min. Then, 1 μM nocodazole was added and images were acquired every 2 min for 1 h. Acquisitions were performed with a Nikon Eclipse Ti-E epifluorescence inverted microscope, equipped with a pco.edge sCMOS camera, Metamorph software and using a 60× 1.49 NA oil objective. Cells were maintained at 5% CO_2_ and 37°C in normal astrocyte medium during acquisition. Scale bar: 20 μm.

***Movie 2: GFP-αTAT1 localisation at FAs upon Y-27632 treatment***. Cells were transfected with GFP-αTAT1 and mCherry-vinculin. First, images were acquired every 2 min for 15 min. Then, 10 μM Y-27632 was added and images were acquired every 2 min for 1 h. Acquisitions were performed with a Nikon Eclipse Ti-E epifluorescence inverted microscope, equipped with a pco.edge sCMOS camera, Metamorph software and using a 60× 1.49 NA oil objective. Cells were maintained at 5% CO_2_ and 37°C in normal astrocyte medium during acquisition. Scale bar: 20 μm.

***Movie 3: Chemical wound setup***. Cells were plated on polyacrylamide hydrogels of different rigidities and grown into a monolayer for 1-2 days. A chemical wound was induced using a microinjector needle, with 0.05 M NaOH. Once the chemical is microdropped on the surface of the monolayer, a circular wound is created. Dead cells and debris are washed out and cells are allowed to migrate for 12 h. Video was acquired using a 10X objective on a Leica DMI 6000B microscope equipped with the Leica software. Cells were maintained at 5% CO_2_ and 37°C in normal astrocyte medium during acquisition.

***Movie 4: Astrocytes migrate faster on stiff substrates***. Wild-type cells were plated on polyacrylamide hydrogels of different rigidities and grown into a monolayer for 1-2 days. After creating a chemical wound, cells were allowed to migrate towards the wound. Images were acquired every 15 min for 12 h using a Zeiss Axiovert 200M, dry objective 10X 0.45 NA and a pco.edge sCMOS camera. Cells were maintained at 5% CO_2_ and 37°C in normal astrocyte medium during acquisition. Scale bar: 25 μm.

***Movie 5: αTAT1 regulates mechanosensitive cell migration***. Cells transfected with siCtl, siαTAT1 were plated on polyacrylamide hydrogels of different rigidities and grown into a monolayer for 2-3 days. After creating a chemical wound, cells were allowed to migrate towards the wound. Images were acquired every 15 min for 12 h using a Zeiss Axiovert 200M, dry objective 10X 0.45 NA and a pco.edge sCMOS camera. Cells were maintained at 5% CO_2_ and 37°C in normal astrocyte medium during acquisition. Scale bar: 25 μm.

## Notes

### Competing Interest Statement

The authors have declared no competing interest.

## References

1 Jaalouk, D. E. & Lammerding, J. Mechanotransduction gone awry. Nature reviews. Molecular cell biology 10, 63–73, doi: 10.1038/nrm2597 (2009).

2 Sun, Z., Guo, S. S. & Fässler, R. Integrin-mediated mechanotransduction. The Journal of cell biology 215, 445–456, doi: 10.1083/jcb.201609037 (2016).

3 Elosegui-Artola, A. et al. Rigidity sensing and adaptation through regulation of integrin types. Nat Mater 13, 631–637 (2014).

4 Ladoux, B., Mège, R.-M. & Trepat, X. Front-Rear Polarization by Mechanical Cues: From Single Cells to Tissues. Trends in Cell Biology 26, 420–433, doi: 10.1016/j.tcb.2016.02.002 (2016).

5 Etienne-Manneville, S. Microtubules in cell migration. Annual review of cell and developmental biology 29, 471–499, doi: 10.1146/annurev-cellbio-101011-155711 (2013).

6 Bouchet, B. P. & Akhmanova, A. Microtubules in 3D cell motility. Journal of cell science 130, 39–50, doi: 10.1242/jcs.189431 (2017).

7 Martins, G. G. & Kolega, J. A role for microtubules in endothelial cell protrusion in three-dimensional matrices. Biology of the cell 104, 271–286 (2012).

8 Seetharaman, S. & Etienne-Manneville, S. Microtubules at focal adhesions – a double-edged sword. Journal of cell science 132, jcs232843, doi: 10.1242/jcs.232843 (2019).

9 Etienne-Manneville, S. & Hall, A. Integrin-mediated activation of Cdc42 controls cell polarity in migrating astrocytes through PKCzeta. Cell 106, 489–498 (2001).

10 Etienne-Manneville, S. In vitro assay of primary astrocyte migration as a tool to study Rho GTPase function in cell polarization. Methods Enzymol 406, 565–578 (2006).

11 Bance, B., Seetharaman, S., Leduc, C., Boëda, B. & Etienne-Manneville, S. Microtubule acetylation but not detyrosination promotes focal adhesion dynamics and astrocyte migration. Journal of cell science 132, jcs.225805, doi: 10.1242/jcs.225805 (2019).

12 Coombes, C. et al. Mechanism of microtubule lumen entry for the α-tubulin acetyltransferase enzyme αTAT1. Proceedings of the National Academy of Sciences 113, E7176–E7184, doi: 10.1073/pnas.1605397113 (2016).

13 Valenzuela-Fernandez, A., Cabrero, J. R., Serrador, J. M. & Sánchez-Madrid, F. HDAC6: a key regulator of cytoskeleton, cell migration and cell–cell interactions. Trends in cell biology 18, 291–297 (2008).

14 Haggarty, S. J., Koeller, K. M., Wong, J. C., Butcher, R. A. & Schreiber, S. L. Multidimensional Chemical Genetic Analysis of Diversity-Oriented Synthesis-Derived Deacetylase Inhibitors Using Cell-Based Assays. Chemistry & Biology 10, 383–396, doi: 10.1016/S1074-5521(03)00095-4 (2003).

15 Haggarty, S. J., Koeller, K. M., Wong, J. C., Grozinger, C. M. & Schreiber, S. L. Domain-selective small-molecule inhibitor of histone deacetylase 6 (HDAC6)-mediated tubulin deacetylation. Proceedings of the National Academy of Sciences 100, 4389–4394, doi: 10.1073/pnas.0430973100 (2003).

16 Choquet, D., Felsenfeld, D. P. & Sheetz, M. P. Extracellular matrix rigidity causes strengthening of integrin-cytoskeleton linkages. Cell 88, 39–48, doi: 10.1016/s0092-8674(00)81856-5 (1997).

17 Wolfenson, H. et al. Tropomyosin controls sarcomere-like contractions for rigidity sensing and suppressing growth on soft matrices. Nature cell biology 18, 33–42, doi: 10.1038/ncb3277 (2016).

18 Peglion, F., Llense, F. & Etienne-Manneville, S. Adherens junction treadmilling during collective migration. Nature cell biology 16, 639 (2014).

19 Janke, C. & Montagnac, G. Causes and Consequences of Microtubule Acetylation. Current Biology 27, R1287–R1292 (2017).

20 Enomoto, T. Microtubule disruption induces the formation of actin stress fibers and focal adhesions in cultured cells: possible involvement of the rho signal cascade. Cell structure and function 21, 317–326 (1996).

21 Bershadsky, A., Chausovsky, A., Becker, E., Lyubimova, A. & Geiger, B. Involvement of microtubules in the control of adhesion-dependent signal transduction. Current Biology 6, 1279–1289, doi: https://doi.org/10.1016/S0960-9822(02)70714-8 (1996).

22 Goult, B. T., Yan, J. & Schwartz, M. A. Talin as a mechanosensitive signaling hub. The Journal of cell biology 217, 3776–3784, doi: 10.1083/jcb.201808061 (2018).

23 Prager-Khoutorsky, M. et al. Fibroblast polarization is a matrix-rigidity-dependent process controlled by focal adhesion mechanosensing. Nature cell biology 13, 1457–1465, doi: 10.1038/ncb2370 (2011).

24 Tran, A. D.-A. et al. HDAC6 deacetylation of tubulin modulates dynamics of cellular adhesions. Journal of cell science 120, 1469–1479, doi: 10.1242/jcs.03431 (2007).

25 De Pascalis, C. et al. Intermediate filaments control collective migration by restricting traction forces and sustaining cell–cell contacts. The Journal of cell biology 217, 3031–3044, doi: 10.1083/jcb.201801162 (2018).

26 Sakamoto, Y., Boeda, B. & Etienne-Manneville, S. APC binds intermediate filaments and is required for their reorganization during cell migration. The Journal of cell biology 200, 249–258, doi: 10.1083/jcb.201206010 (2013).

27 Schiller, H. B. et al. beta1- and alphav-class integrins cooperate to regulate myosin II during rigidity sensing of fibronectin-based microenvironments. Nature cell biology 15, 625–636, doi: 10.1038/ncb2747 (2013).

28 Lawson, C. D. & Ridley, A. J. Rho GTPase signaling complexes in cell migration and invasion. 217, 447–457, doi: 10.1083/jcb.201612069 (2018).

29 Krendel, M., Zenke, F. T. & Bokoch, G. M. Nucleotide exchange factor GEF-H1 mediates cross-talk between microtubules and the actin cytoskeleton. Nature cell biology 4, 294–301, doi: 10.1038/ncb773 (2002).

30 Ren, Y., Li, R., Zheng, Y. & Busch, H. Cloning and characterization of GEF-H1, a microtubule-associated guanine nucleotide exchange factor for Rac and Rho GTPases. The Journal of biological chemistry 273, 34954–34960 (1998).

31 Heck, J. N. et al. Microtubules regulate GEF-H1 in response to extracellular matrix stiffness. Molecular biology of the cell 23, 2583–2592, doi: 10.1091/mbc.E11-10-0876 (2012).

32 Rafiq, N. B. M. et al. A mechano-signalling network linking microtubules, myosin IIA filaments and integrin-based adhesions. Nature Materials 18, 638–649, doi: 10.1038/s41563-019-0371-y (2019).

33 Jiu, Y. et al. Vimentin intermediate filaments control actin stress fiber assembly through GEF-H1 and RhoA. 130, 892–902, doi: 10.1242/jcs.196881 (2017).

34 Trepat, X. et al. Physical forces during collective cell migration. Nature physics 5, 426 (2009).

35 Fernández-Barrera, J. et al. The actin-MRTF-SRF transcriptional circuit controls tubulin acetylation via <em>α-TAT1</em> gene expression. The Journal of cell biology 217, 929–944, doi: 10.1083/jcb.201702157 (2018).

36 Thurston, S. F., Kulacz, W. A., Shaikh, S., Lee, J. M. & Copeland, J. W. The ability to induce microtubule acetylation is a general feature of formin proteins. PloS one 7, e48041, doi: 10.1371/journal.pone.0048041 (2012).

37 Portran, D., Schaedel, L., Xu, Z., Thery, M. & Nachury, M. V. Tubulin acetylation protects long-lived microtubules against mechanical ageing. 19, 391–398, doi: 10.1038/ncb3481 (2017).

38 Xu, Z. et al. Microtubules acquire resistance from mechanical breakage through intralumenal acetylation. Science (New York, N.Y.) 356, 328–332, doi: 10.1126/science.aai8764 (2017).

39 Barriga, E. H., Franze, K., Charras, G. & Mayor, R. Tissue stiffening coordinates morphogenesis by triggering collective cell migration in vivo. Nature 554, 523–527, doi: 10.1038/nature25742 (2018).

40 Sunyer, R. et al. Collective cell durotaxis emerges from long-range intercellular force transmission. Science (New York, N.Y.) 353, 1157–1161, doi: 10.1126/science.aaf7119 (2016).

41 Serra-Picamal, X., Conte, V., Sunyer, R., Munoz, J. J. & Trepat, X. Mapping forces and kinematics during collective cell migration. Methods in cell biology 125, 309–330, doi: 10.1016/bs.mcb.2014.11.003 (2015).

42 Tambe, D. T. et al. Collective cell guidance by cooperative intercellular forces. Nature materials 10, 469 (2011).

43 Heuser, J. The production of ‘cell cortices’ for light and electron microscopy. Traffic (Copenhagen, Denmark) 1, 545–552, doi: 10.1034/j.1600-0854.2000.010704.x (2000).

44 Martiel, J. L. et al. Measurement of cell traction forces with ImageJ. Methods in cell biology 125, 269–287, doi: 10.1016/bs.mcb.2014.10.008 (2015).

45 Poullet, P., Carpentier, S. & Barillot, E. myProMS, a web server for management and validation of mass spectrometry-based proteomic data. Proteomics 7, 2553–2556, doi: 10.1002/pmic.200600784 (2007).

46 Valot, B., Langella, O., Nano, E. & Zivy, M. MassChroQ: a versatile tool for mass spectrometry quantification. Proteomics 11, 3572–3577, doi: 10.1002/pmic.201100120 (2011).

47 Vizcaino, J. A. et al. 2016 update of the PRIDE database and its related tools. Nucleic acids research 44, D447–456, doi: 10.1093/nar/gkv1145 (2016).

48 Perez-Riverol, Y. et al. The PRIDE database and related tools and resources in 2019: improving support for quantification data. Nucleic acids research 47, D442–d450, doi: 10.1093/nar/gky1106 (2019).

